# Glycan analysis of human neutrophil granules implicates a maturation-dependent glycosylation machinery

**DOI:** 10.1101/2020.04.02.021394

**Authors:** Vignesh Venkatakrishnan, Regis Dieckmann, Ian Loke, Harry Tjondro, Sayantani Chatterjee, Johan Bylund, Morten Thaysen-Andersen, Niclas G. Karlsson, Anna Karlsson-Bengtsson

**Author notes:** These authors contributed equally (Vignesh Venkatakrishnan, Regis Dieckmann). **Corresponding author** Dr. Vignesh Venkatakrishnan, Department of Rheumatology and Inflammation Research, Institute of Medicine, Sahlgrenska Academy, University of Gothenburg, Guldhedsgatan 10A, 413 46 Gothenburg, Sweden.

## Abstract

Protein glycosylation is essential to trafficking and immune functions of human neutrophils. During granulopoeisis in the bone marrow, distinct neutrophil granules are successively formed. Distinct receptors and effector proteins, many of which are glycosylated, are targeted to each type of granule according to their time of expression, a process called ‘targeting-by-timing’. Therefore, these granules are time capsules reflecting different times of maturation that can be used to understand how glycosylation evolves during granulopoiesis. Herein, neutrophil subcellular granules were fractionated by Percoll density gradient centrifugation and *N*- and *O*-glycans present in each compartment were analyzed by liquid chromatography and tandem mass spectrometry. We found abundant paucimannosidic *N*-glycans and lack of *O*-glycans in early-formed azurophil granules (AG), whereas later-formed specific and gelatinase granules (SG and GG) contained complex *N*- and *O*-glycans with remarkably elongated *N*-acetyllactosamine repeats with Lewis-x and sialyl-Lewis-x epitopes. Many glycans identified are unique to neutrophils and their complexity increased progressively from AG to SG and then to GG, suggesting temporal changes in the glycosylation machinery indicative of ‘glycosylation-by-timing’ during granulopoiesis. In summary, this comprehensive neutrophil granule glycome map, the first of its kind, highlights novel granule-specific glycosylation features and is a crucial first step towards a better understanding of the mechanisms regulating protein glycosylation during neutrophil granulopoiesis and a more detailed understanding of neutrophil biology and function.

## Introduction

Neutrophils are central cells of innate immunity, primarily dedicated to the killing of invading microbes. They are non-dividing, short-lived white blood cells that, in large numbers and with impressive specificity, can perform numerous functions. It is well-established that mature neutrophils in circulation contain different granule subsets that harbor distinct proteins and other biomolecules ^1,2^. Sequential mobilization of granules allows for the cells to rapidly change their surface receptor repertoire and release matrix-destroying proteases to facilitate transmigration from the blood, extravasation into infected and inflamed tissues, and phagocytosis of microbes and cellular debris. Therefore, a distinct separation of neutrophil proteins between the different granules and vesicles is of importance for the ability to perform appropriately in response to inflammation and infection.

The neutrophil granules are sequentially formed during granulopoiesis, the process of neutrophil maturation in the bone marrow. Early during the promyelocyte stage, the azurophil granules (AG) are formed, followed by the specific and gelatinase granules (SG and GG) which develop sequentially during the myelocyte/metamyelocyte and band cell stages^1^. Finally, the so-called secretory vesicles (SV) are formed from the plasma membrane (PM) of mature neutrophils through endocytosis. Since no subsequent exchange of proteins between fully synthesized compartments are thought to occur, the neutrophil proteins that are synthesized during different stages of granulopoiesis end up in the corresponding granule type, a process known as ‘targeting-by-timing’. This was elegantly exemplified in studies utilizing proteomic analyses of neutrophil granules, SV and PM ^1,3,4^.

The majority of granule and cell surface proteins are glycosylated^1^. Glycosylation adds structural and functional heterogeneity to proteins by the attachment of oligosaccharides to specific amino acid residues. *N*-glycans are attached to amino acid consensus sequences Asn-x-Ser/Thr (x ≠ Pro) while *O*-glycans are attached to Ser/Thr/Tyr. Specific glycan structures and/or epitopes are involved in multiple aspects of the immune response, including neutrophil function. The glycosylation of neutrophil proteins is diverse and functionally important for the inflammatory response, e.g., by binding to immune regulating lectins such as selectins, siglecs, galectins and microbial adhesins ^5-9^.

Earlier mass spectrometry (MS) studies documenting protein glycosylation in whole intact neutrophils showed the presence of Galβ1-4GlcNAc, i.e., *N*-acetyl-lactosamine chains (LacNAc) linked to lipids or protein ^10,11^ and that some of the LacNAc residues were modified with fucose and/or terminal sialic acid, generating epitopes such as Lewis x (Le^x^; Galβ1-4(Fucα1-3)GlcNAcβ-R) and sialyl-Lewis x (sLe^x^; Neu5Acα2-3Galβ1-4(Fucα1-3)GlcNAcβ-R). These epitopes are the main ligands of endothelial selectins involved in neutrophil capture and rolling on the endothelium, enabling transmigration to the tissue ^12,13^. Deficiency in the sLe^x^ antigen at the cell surface causes leukocyte adhesion deficiency 2 (LAD2), associated with recurrent infection, persistent leukocytosis and severe mental and growth retardation ^14,15^. The interest in the roles played by Le^x^ and sLe^x^ antigens in cell recruitment has resulted in more specific analyses of these epitopes in neutrophils, on a subcellular level.

Detailed analyses of glycan structures of certain neutrophil granule glycoproteins have been documented ^9,16-18^. Although glycosylation is essential to neutrophil function, a comprehensive mapping of the granule and cell surface glycome is still lacking. Recently, we have identified paucimannosidic glycans (Man_1-3_GlcNAc_2_Fuc_0-1_) carried by a range of neutrophil glycoproteins in the sputum of cystic fibrosis patients ^19^. These glycoproteins were traced back to the AG of human neutrophils ^19,20^, for example, neutrophil elastase, where the unusually truncated glycans were shown to influence the binding of elastase to mannose-binding lectin and α1-anti-trypsin often present within inflamed tissues ^21^.

Herein, we provide a comprehensive characterization of the *N*- and *O*-glycans in neutrophil granules, isolated using subcellular fractionation and analyzed by MS. We show vast differences in glycosylation between different granules. By building on the ‘targeting-by-timing’ hypothesis, the granule specific glycan differences therefore suggests that the glycosylation machinery undergoes alterations during granulopoiesis. The differences in glycosylation between granules can impact immune modulating functions of neutrophils during inflammation.

## Materials and Methods

### Isolation of human neutrophils and subcellular fractionation

Human peripheral blood polymorphonuclear neutrophils (PMNs) were isolated from buffy coats (The Blood Center, Sahlgrenska University Hospital, Gothenburg) obtained from healthy donors ^22^. The erythrocytes were removed by dextran (1%) sedimentation (1 ×g). Monocytes and lymphocytes were removed by centrifugation on a Ficoll–Paque density gradient. The cells were then washed in Krebs-Ringer phosphate buffer (KRG) containing glucose (10 mM) and Mg^2+^ (1.5 mM) and remaining erythrocytes were removed by hypotonic lysis. These purified neutrophils (> 95% pure) were fractionated according to Borregaard et al.^*23*^. In short, 10^9^ cells were treated with the serine protease inhibitor diisopropylfluorophosphate (DFP, final concentration 8 mM), resuspended in 10 mL homogenization buffer [100 mM KCl, 3 mM NaCl, 3.5 mM MgCl_2_, 10 mM PIPES,; pH 7.4] with 1 mM ATP(Na)_2_, 0.5 mM PMSF, and disrupted in a nitrogen bomb (Parr Instruments, Moline, IL, USA) at 400 psi (2500 kPa). Intact cells and nuclei were removed by successive centrifugation steps at 300 x g and 800 x g, respectively. The post nuclear supernatant was layered on top of a two-layer (1.05 and 1.12 g/L) or a three-layer (1.05, 1.09 and 1.12 g/L) Percoll gradient and centrifuged at 15 000 x g for 45 min in a fixed-angle JA-20 Beckman rotor. The material was typically collected in 25 fractions of 1.5 mL (two-layer gradients) or 32 fractions of 1.0 mL (three-layer gradients) from the bottom of the centrifuge tube.

### Granule purity and fractionation

The distribution of four different, established markers (myeloperoxidase [MPO], marker for the AG; lactoferrin [LF], marker for the SG; matrix metalloproteinase-9 [MMP9], marker for the GG; and alkaline phosphatase [ALP], marker for the SV and plasma membrane [PM]) were determined to establish the distribution of the different organelles in the gradient. Enzyme activity of MPO and alkaline phosphatase, respectively, was determined as previously described by Feuk-Lagerstedt E et al. ^24^. The LF and MMP9 distribution was assessed by immunoblot using antibodies (anti-LF, Dako, rabbit polyclonal, 1/2500; anti-MMP9, Calbiochem, rabbit polyclonal n°444236, 1/2000). When appropriate, fractions containing the different organelles were pooled, diluted in homogenization buffer and ultracentrifugated (100,000 x g, 1H) to concentrate the sample and to remove the Percoll. To separate the granule luminal content from the granule membranes, granule pellets were diluted in homogenization buffer, disrupted with a tip sonicator at 1/3 power (8 MHz) for 3 × 30 seconds on ice and fractionated by ultracentrifugation (100’000 x g, 1H, Beckman TLA 120.2 rotor). The luminal fraction was concentrated to around 50 µL on a 3 kDa MWCO Amicon Ultra 0.5 mL centrifugal filter while membrane pellets were dissolved in 50 µL TBS with 2% SDS. For subsequent glycome analyses, protein concentration of total, luminal and membrane fractions of granules was determined by BCA assay (ThermoScientific) using bovine serum albumin as standard.

### Detection of Lewis antigens by immunoblot and immunofluorescence

The primary antibodies used were directed against Le^x^ (mouse monoclonal, clone HI98, Santa Cruz Biotechnology) and sLe^x^ (mouse monoclonal, clone CSLEX-1, BD Biosciences). For the detection of Lewis antigens by immunoblots, pooled granule fractions were blotted onto a PVDF membrane, essentially as described ^24^. The membrane was blocked 1 hour at room temperature (RT) in TBS, 0.1% BSA. Primary antibodies were diluted 1/500 in TBS, 0.01% Tween 20 and incubated at 4°C, overnight (O/N). Immunodetection was performed using secondary horseradish peroxidase-coupled rabbit anti-mouse (P0260, Dako, Denmark) with Clarity Western ECL Substrate (170-5061, BioRad). For the detection of Lewis antigens by immunofluorescence, isolated PMNs were resuspended in cold PBS at 1.5 million cells/mL and let attach to adhesion slides (Marienfeld, Germany) according to the manufacturer’s instructions. Cells were then fixed in 3.5% PFA in PBS and permeabilized with 1% Triton X-100, 10 min. After extensive washing, blocking was performed 30 min, in 0.1% BSA in PBS. Primary antibodies were diluted 1/50 in 0.1% BSA in PBS and incubated 1 hour. Immunodetection was performed using secondary goat anti-mouse IgM Alexa 488 (A-11001, ThermoFisher Scientific), diluted 1/300 in 0.1% BSA in PBS and incubated 1 hour. Imaging was performed with a confocal laser scanning LSM-700 microscope (Zeiss).

### Release of *N*-glycans from neutrophil granule proteins

The isolated neutrophil granule proteins were solubilized in 6.0 M urea and 0.1% SDS. *N*- and *O*-glycans from the solubilized proteins were released and processed as described here ^25^. In brief, 100 µg protein were dot blotted on PVDF membrane (Millipore), stained with Alcian blue and excised. *N*-glycans were released from the intact proteins using 0.5 U/ µL *N*-glycosidase F (PNGase F, *Elizabethkingia miricola*, Promega), overnight, at 37°C. The released *N*-glycans were reduced to alditols by treatment with 0.50 M NaBH4 in 50mM KOH for 3 h at 50°C. The reduction was quenched by addition of 2 µL of glacial acetic acid. The reduced *N*-glycans were desalted on a strong cation exchange micro-column on top of hydrophobic C18 material. The *N*-glycans were further cleaned up with methanol to eliminate the presence of boron, dried by vacuum centrifugation and re-dissolved in 10 µL of water for LC-MS/MS analysis.

Purified neutrophil granule proteins, both from luminal and membrane fractions (∼20 µg) were solubilized in homogenization buffer and reduced using 10 mM dithiothreitol (DTT) in 100 mM ammonium bicarbonate (pH 8.4), 45 min, 56°C and alkylated using 25 mM iodoacetamide in 100 mM ammonium bicarbonate (pH 8.4) (both final concentration), 30 min in the dark, 22°C. Proteins was then blotted on a primed 0.45 µm PVDF membrane and stained with Direct Blue. Stained protein spots were excised, transferred to separate wells in a flat bottom polypropylene 96 well plate, blocked with 1% w/v polyvinylpyrrolidone in 50%v/v methanol and washed with MillQ water. N-glycans were enzymatic released using 3.5 U Flavobacterium meningosepticum N-glycosidase F (Promega) in 20µl water/well, 16 h, 37 °C. The unstable amino group (-NH_2_) of the reducing and GlcNAC residues of N-glycosidase F-released N-glycans were allowed to spontaneously convert to hydroxyl groups (-OH) in weak acid using 100 mM ammonium acetate, pH 5, 1 h, 22°C to facilitate subsequent quantitative reduction to glycan alditols. Reduction was carried out using 1 M sodium borohydride in 50 mM potassium hydroxide, 3 h, 50°C. The reaction was stopped by glacial acetic acid quenching. Dual desalting was then performed in micro-sold phase extraction (SPE) formats using strong cation exchange/C18 (where N-glycans are not retained) and porous gaphitised carbon (PGC) (where N-glycans are retained) as stationary phases, respectively. The desalted N-glycans were eluted from the PGC-SPE columns using 40% v/v ACN containing 0.1% v/v aqueous TFA, dired and dissolved in 10 µl MilliQ water for N-glycan analysis. Bovine fetuin carrying sialo-N-glycans were included as control glycoproteins to ensure efficient N-glycan release, clean-up and analysis.

### PGC-LC-MS/MS based *N*-glycomics

Released *N*-glycans were analyzed by liquid-chromatography-tandem mass spectrometry (LC-MS/MS) in two different laboratories. Overall granule glycans were analyzed on porous graphitized carbon (PGC)-LC using a 10 cm × 250 µm i.d. column (in-house), containing 5 µm PGC particles (Thermo Scientific, Waltham, MA, USA) connected to an LTQ-ion trap mass spectrometer (Thermo Scientific). The *N*- and *O*-glycans were eluted using a linear gradient from 0 to 40% acetronitrile in 10 mM ammonium bicarbonate over 40 min at a flow rate of 250 µL/min split down to 5-10 µL/min. Electrospray ionization-mass spectrometry (ESI-MS) was performed in negative ion polarity with an electrospray voltage of 3.5 kV, capillary voltage of −33.0 V, and capillary temperature of 300°C. The following scan events were used: MS full scan (*m/z* 380-2000) and data-dependent tandem MS (MS/MS) scans after collision-induced dissociation (CID) on precursor ions at a normalized collisional energy of 35% with a minimum signal of 300 counts, isolated width of 2.0 *m/z*, and activation time of 30 ms. The data were viewed and manually analyzed using Xcalibur software (version 2.2, Thermo Scientific).

*N*-glycans released from each granule membrane and soluble proteins were analyzed using capillary LC-MS/MS (Dionex Ultimate 3000) on an ESI linear ion trap mass spectrometer (LTQ Velos Pro, Thermo Scientific, Melbourne, Australia). Samples were injected onto a PGC-LC capillary column (Hypercarb KAPPA, 5 μm particle size, 200 Å pore size, 180 μm inner diameter × 100 mm length, Thermo Scientific) kept at 50°C. Separation of N-glycans was carried out over a linear gradient of 2–32% ACN/10 mm ammonium bicarbonate for 75 min at a constant flow rate of 4 μl min−1. The sample injection volume was 4 μl. The acquisition range was m/z 380–2000. The acquisition was performed in negative ionisation polarity with an electrospray voltage of 4.0 kV, capillary voltage of −14.9 kV and capillary temperature of 310°C. Data-dependent acquisition were used where the top nine most abundant precursors in each full scan spectrum were selected for MS/MS using collision induced dissociation (CID) at a normalised collision energy of 35%. A minimum signal of 300 counts was required for MS/MS. The precursor isolation width was m/z 1.0 and the activation time was 10 msec. The mass spectrometer was calibrated using a tune mix (Thermo Scientific). Mass spectra were viewed and analysed using Xcalibur V2.2 (Thermo Scientific). Glycoworkbench assisted in the annotation and visualisation of the N-glycan structures ^26^.

### Structural determination of *N*-glycans and relative quantitation

The monosaccharide compositions and each *N*-glycan structural sequence and linkage were manually analyzed using monoisotopic mass, CID-MS/MS fragmentation and absolute retention time on the PGC-LC-MS/MS N-glycome data. Monoisotopic molecular masses of the identified glycans, which were generally within 0.5 Da of the theoretical masses, were matched against likely mammalian *N*-glycan monosaccharide compositions using GlycoMod web tool. GlycoWorkBench v2.1 was used to draw the proposed *N*-glycans and to generate *in silico* glycosidic and cross-ring fragments to assist in the structural interpretation.

### Structural assumptions

The linkages for chitobiose core, the initial arm extension including β1,2 GlcNAc-Man linkages on the 3’ and 6’ arm for mono- and biantennary structures, β1,4 linkage for Gal-GlcNAc and α1,6 link for core fucosylation were assumed from the established knowledge of human *N*-glycosylation biosynthetic machinery. Substantial in-house knowledge of the absolute and relative PGC retention behavior of mammalian *N*-glycans, in particular for isomeric linkage-type *N*-glycans, was used to increase the confidence of the reported structures. Structural ambiguity, in particular for terminal monosaccharide, including fucosylation and sialylation, remains unresolved for some structures, where knowledge of the structural relation with PGC retention time and/ or confirming CID fragmentation were absent.

### Statistical analysis

Statistical analysis was performed using GraphPad Prism version 8 software (La Jolla, CA). The quantitative data are expressed as mean ± SEM and analyzed using two-tailed Student’s *t*-test and *p* values ≤ 0.05 were considered statistically significant.

## Results

### The neutrophil subcellular organelles are characterized by unique *N*-glycan signatures

Subcellular organelles were first separated by two-layer gradients into three fractions; AG, SG+GG, and SV+PM (Figure 1a). Solubilization of organelle proteins was followed by enzymatic release of *N*-glycans and MS analyses enabled the unambiguous identification of 71 unique *N*-glycans. The obtained profiles between the three fractions differed vastly from each other (Figure 1b). Paucimannosidic and complex *N*-glycans were predominantly identified in AG and SG+GG, respectively, whereas SV+PM contained oligomannose and complex glycans, at similar levels (40-50%) (Figure 1b). The analysis identified 36 *N*-glycans in AG, 60 in SG+GG, and 61 in the SV+PM. Out of the total 71 *N*-glycans, 28 were present in all organelles and SG+GG and SV+PM were mostly similar, sharing 53 *N*-glycans (Figure 1c). Qualitative differences were corroborated also on the quantitative level (Figure 1d). Hence, the different organelles all had unique sets of glycan structures that were in some parts overlapping. The detailed list of *N*-glycans and their relative abundances in each fraction are found in supplementary table.

**Figure 1.**
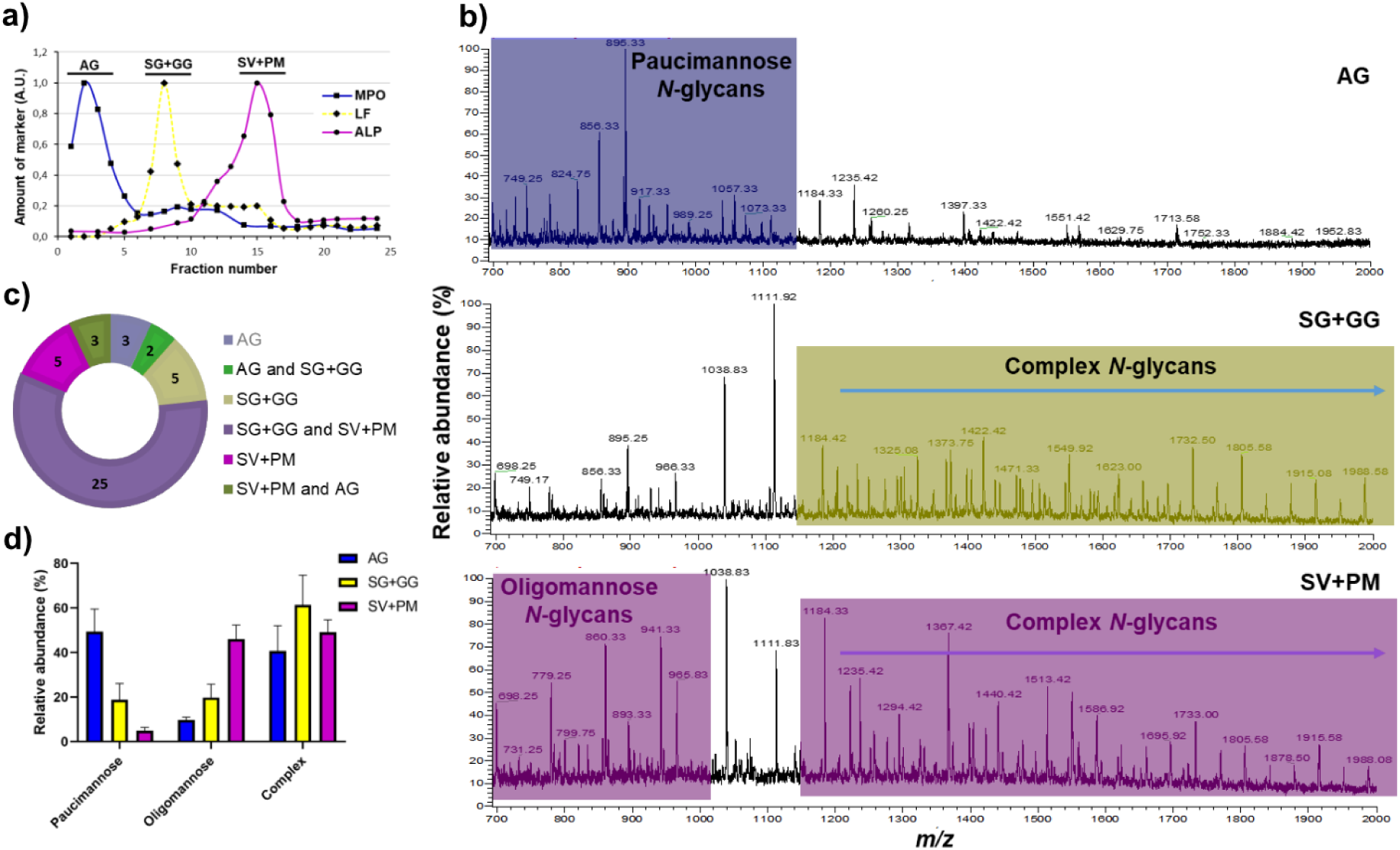
Validation of neutrophil subcellular fractionation and organelle-specific neutrophil *N*-glycan profiling. a) The granules of naive neutrophils were fractionated using a two-layer Percoll gradient. Fractions were assigned to distinct neutrophil organelles based on the level of known granule-specific protein markers as determined using immunoblot and activity assays (y-axis, arbitrary units, AU): MPO activity (marker for azurophil granules, AG, filled squares); lactoferrin (marker for specific granules, SG, filled diamonds); total alkaline phosphatase (ALP) activity (measured in the presence of detergent; marker for secretory vesicles and plasma membrane, SV+PM, filled circles). Data shown are representative from three separate, independent experiments. b) Mass spectral profiles of *N*-glycans released from neutrophil granule protein extracts, showing dramatic qualitative *N*-glycan profile differences between the azurophil granules (AG), the specific and gelatinase granules (SG+GG), and secretory vesicles and plasma membrane (SV+PM) fractions. Characteristic *N*-glycan types for each granule fraction have been color coded to highlight the most prominent differences. c) Distribution of the identified *N*-glycan structures in the different neutrophil subcellular compartments. d) The *N*-glycans identified in the different subcellular fractions were grouped into the major *N*-glycan types i.e. paucimannosidic, oligomannosidic and complex *N*-glycans and their relative abundance were determined for each granule subset. Data points are plotted as the mean ± SEM (n = 3).

### Paucimannosidic and oligomannoseidic-type *N*-glycans are abundant in AG and SV+PM, respectively

The hallmark of AG was the presence of paucimannosidic glycans, with a relative abundance of 49.4% ± 5.8 of all *N*-glycans. In contrast, only 18.7% ± 5.2 and 4.9% ± 0.9, respectively, of all *N*-glycans were paucimannosidic in SG+GG and SV+PM (Figure 1d). Six different paucimannosidic glycans were identified in AG, with the most abundant (65% of the paucimanose glycans) containing two mannoses and a core fucose (M2F; *m/z* 895.3^1-^) (Figure 2a, Supplementary figure S1). In fact, within the class of paucimannosidic-type N-glycans, the M2F was clearly the most abundant paucimannosidic glycan species in all organelles (Figure 2a), but the overall abundance of M2F in SV+PM and SG+GG was <5% of total *N*- glycans as compared to 40% in the AG Another striking feature of AG is the high abundance of core fucosylation (80.4%) as compared to terminal fucosylation (19.6%).

**Figure 2.**
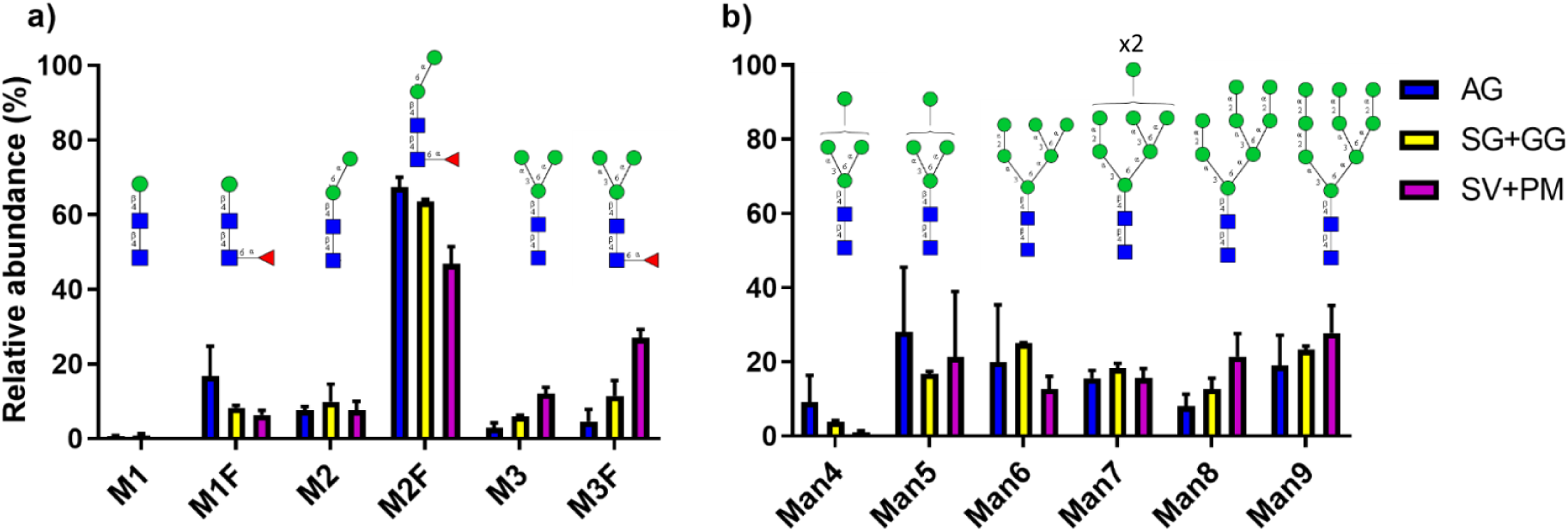
Distribution of mannose-terminating *N*-glycans in AG and SV+PM. Relative abundance of the observed a) paucimannosidic *N*-glycans and b) oligomannose *N*-glycans in AG, SG+GG and SV+PM fractions. The relative distribution of *N*-glycans are plotted as the mean ± SEM, n = 3. Blue square, GlcNAc; green circle, mannose; red triangle, fucose

Oligomannose and complex type *N*-glycans predominates in the SV+PM fraction, with a relative abundance of 46.1% ± 3.7 and 49.1% ± 3.2, respectively (Figure 1b, 1d). Oligomannose *N*-glycans in the SV+PM fraction showed low abundance of Man4 glycan and a varied distribution of Man5 to Man9 glycans (Figure 2b). Oligomannose glycans were present also in AG and SG+GG, however at much lower levels (9.7% ± 0.7 and 19.9% ± 4.2, respectively). The distribution between Man4 to Man9 glycans was similar in these organelles as in SV+PM **(Figure 2b)**.

### Complex *N*-glycans with LacNAc repeats are characteristic of the SG+GG and SV+PM

Complex *N*-glycans were observed in all fractions, with a higher abundance in the SG+GG (61.4% ± 9.5 of all *N*-glycans) as compared to the AG (40.7% ± 6.5) and SV+PM (49.1% ± 3.2) (Figure 1d). Conventional glycans carrying single LacNAcs on both antennae were observed in AG, albeit in lower abundance (5.8% ± 1.8) as compared to the other fractions. In contrast, one of the key glycosylation features in SG+GG and SV+PM was the presence of unusual, very high molecular mass complex glycans extended with repeating LacNAc units (Figure 3a). As these glycans were characterized by broad peaks of extracted precursor ions due to unresolved multiple isomers their abundances were not considered during our initial analysis.

**Figure 3.**
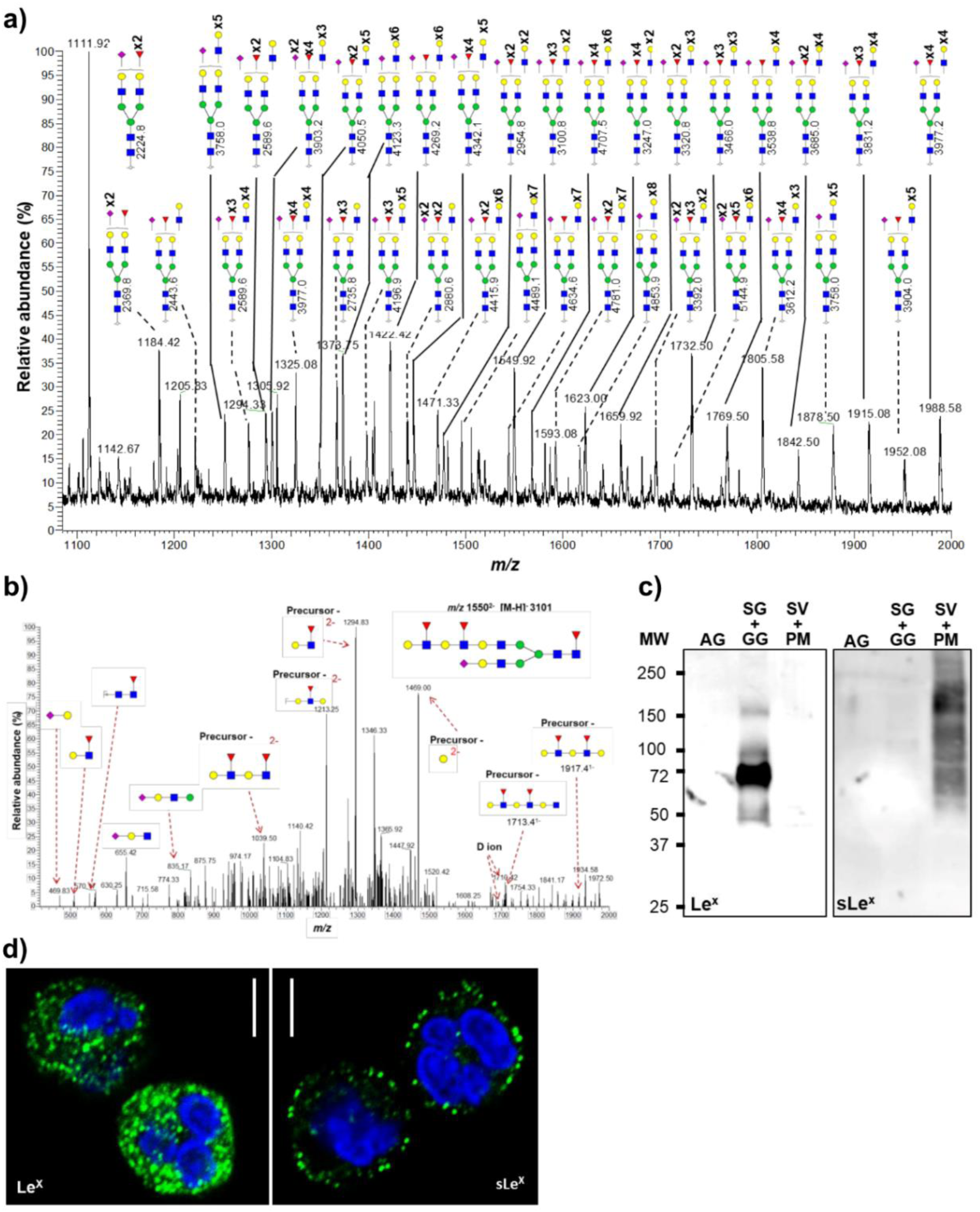
Unusual LacNAc-elongated complex *N*-glycans reside in the specific and gelatinase granules. a) Representative mass spectral profile of SG+GG *N*-glycans illustrating the presence of unusual LacNAc-elongated *N*-glycans, up to 10 repeats, with and without fucose and/or sialic acid capping, forming Lewis epitopes. b) *N*-glycans were characterized using CID-MS/MS (M-H) as exemplified by the annotated MS/MS fragmentation spectrum of the precursor ion *m/z* 1550^2-^ corresponding to an *N*-glycan containing four LacNAc units with fucosyl and sialyl decorations. Diagnostic ions represents the presence of LacNAc elongation as opposed to antenna branching are shown here. c) Immunoblot analysis of Le^x^ and sLe^x^ epitopes on proteins from different organelles. Equal protein amounts were loaded for each fraction as determined by BCA assay. d) Representative images of immunofluorescence-stained neutrophils demonstrating the localization of Le^x^ (left image) and sLe^x^ (right image) epitopes. Scale bar = 5 µm. Blue square, *N*-acetyl glucosamine; green circle, mannose; red triangle, fucose; yellow circle, galactose; purple diamond, sialic acid (*N*-acetyl neuraminic acid).

Apart from oligomannose structures, the glycan profile of SV+PM was similar to that of the SG+GG with elongated complex *N*-glycans (Supplementary Figure S2). In SG+GG, complex-type *N*-glycans were predominantly bi-antennary fucosylated glycans with or without terminal sialic acid. The most abundant glycan (*m/z* 1111.9^2-^) identified in SG+GG was a bi-antennary mono-sialylated and di-fucosylated glycan with core fucosylation and terminal fucosylation observed on a single antenna (Figure 3a). In conclusion, complex *N*-glycans with LacNAc repeats were abundant in SG+GG and SV+PM, but absent from AG.

### The LacNAc repeats are mainly found on biantennary *N*-glycans

The structures of the extended LacNAcs were confirmed based on their composition related to their molecular mass, along with supporting spectral data obtained through MS/MS fragmentation. The compositions were found to be consistent with extended versions of accurately assigned lower molecular mass *N*-glycans with LacNAc repeats. A representative MS/MS fragmentation of a four-LacNAc containing glycan with an (*m/z* 1550^2-^) is shown in Figure 3b. The presence of various fragment ions, including *m/z* 1039^2-^, 1213^2-^, 1294^2-^, 1469^2-^, 1713^1-^, 1917^1-^ and D-ions of 1692^1-^ and 1710^1-^ suggests i) elongation of 6’ arm rather than branched tri- or tetra-antennary glycans and ii) LacNAc extension on a single arm. These fragment ions were observed in the MS/MS fragmentation spectrum of other glycans as well (Supplementary Figure S3), which supports and confirms our assessment. Furthermore, the monosaccharide composition of glycans with additional LacNAc repeats contains either one or two terminal sialic acids, supporting the presence of elongated bi-antennary glycans but not branched glycans. Broad signal peaks of higher *m/z* values suggest that tri- and tetra-antennary glycans elongated with LacNAc units may also be present but at a lower abundance.

Information about the degree of LacNAc extensions could be obtained from the combined intensities of the broad LC-peaks which corresponds to structures potentially carrying three to ten LacNAc units (Supplementary Figure S4). It could be seen that structures containing six to eight LacNAc units were similarly most abundant in SG+GG and SV+PM. Simple extrapolation of the data suggests that even longer than 10 LacNAc unit structures could be present in both these fractions.

### The unusually long LacNAc repeats of SG+GG and SV+PM are decorated with Lewis-like epitopes

Fucosylation of the GlcNAc moiety in the LacNAc chains, building Lewis type epitopes, was observed in the SG+GG and SV+PM fractions. It should be noted that not all LacNAcs in an antenna were fucosylated. For high molecular weight glycans we calculated the number of Lewis epitopes per structure and their corresponding relative intensities (Supplementary Figure S4). Simple structures containing one to three Lewis epitopes were more abundant than structures containing higher numbers of fucose. Structures were found to be decorated with a maximum of seven fucose residues. The data also show that the amount of fucosylation per LacNAc was constantly decreasing with increasing LacNAc repeats.

When analyzing the fractions for Le epitopes by immunoblotting Le^x^ antigens were mainly found in the SG+GG fraction, whereas sLe^x^ epitopes were detected predominantly in the SV+PM fraction (Figure 3c). Using confocal microscopy, the presence of Le^x^ was detected in intracellular granules, whereas sLe^x^ was mainly localized to vesicles close to the cytoplasmic membrane, suggesting that the latter structures are associated with the SV and PM compartments (Figure 3d). Overall, data show that neutrophils contain enormous amounts of complex *N*-glycans with elongated LacNAc repeats that are decorated with fucose (Le^x^; in SG+GG), and capped with sialic acid (sLe^x^; in SV+PM).

### Gelatinase granules carry *N*-glycans with higher numbers of LacNac repeats than specific granules

So far, it is clear that the neutrophil granules, in addition to their difference in size, buoyancy and protein composition, also differs in their glycan composition (Figure 1b and 1d) ^3,27^. Thus, we wanted to investigate whether this is true also for their respective glycan content. The SG and GG fractions were separated from each other using three-layer Percoll gradients (Figure 4a) and analyzed for high molecular weight *N*-glycans.

**Figure 4.**
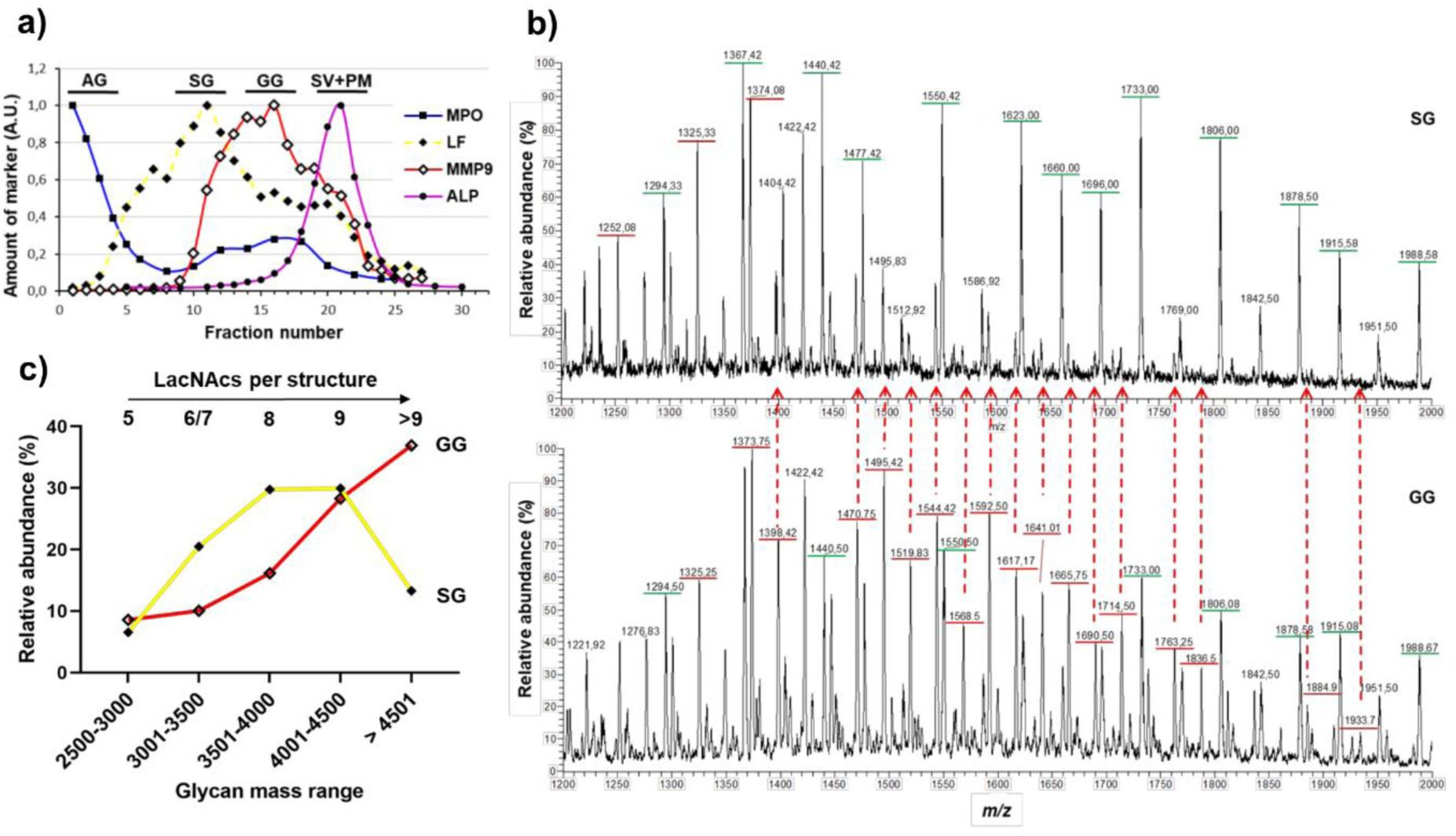
The elongation length of LacNAc-containing *N*-glycans differs in the specific and gelatinase granules. a) The granules of naive neutrophils were fractionated using a three-layer Percoll gradient. Fractions were assigned to distinct neutrophil granule fractions based on the level of known granule-specific protein markers as determined using immunoblots and activity assays (y-axis, arbitrary units, AU): MPO activity (marker for azurophil granules, AG, filled squares); lactoferrin (marker for specific granules, SG, filled diamonds); MMP9 (marker for gelatinase granules, GG, open diamonds); total alkaline phosphatase (ALP) activity (measured in the presence of detergent; marker for secretory vesicles and plasma membrane, SV+PM, filled circles). Data is representative from three separate, independent experiments. The bars indicate which fraction numbers were pooled together for the glycan analysis. b) Representative mass spectra focusing on the high *m/z* region encompassing the elongated LacNAc repeat-containing *N*-glycans released from SG (top) and GG (bottom) protein extracts, respectively. The *m/z* underlined in green and red are doubly and triply charged ions, respectively. Many triply charged glycans that are found in the GG spectrum are either low in abundance or absent in the SG spectrum, indicated by the red dashed arrows. c) The relative intensities of glycan structures identified in SG and GG (based on predicted monosaccharide compositions) and divided into different mass ranges, showing that the GG have more glycans of higher mass than the SG. The number of potential LacNAcs in each mass range is shown.

Mass spectra of *N*-linked glycans in the *m/z* 1200-2000 region revealed an enrichment of triply charged ions in the GG while doubly charged ions were more abundant in the SG (Figure 4b). It is important to note that the detection of more high molecular mass glycans in the isolated GG fraction (three-layer gradient) as compared to in the combined SG+GG fraction (two-layer gradient) is a consequence of absence of SG in the GG fraction. Consequently, the precursor ion (*m/z* 1933.7^3-^) corresponding to 11 LacNAcs, four fucoses and one sialic acid (Δ*mass* of − 0.016), the longest glycan detected, was identified only in the GG fraction of the three-layer gradient.

The relative intensities of glycans, based on their predicted monosaccharide compositions, were grouped into different mass ranges. It is evident that the relative intensities of glycans containing five to eight LacNacs per structure were more abundant in SG as compared to GG (Figure 4c). In contrast, structures containing nine or more LacNAcs were more abundant in the GG fraction. Taken together, the data strongly suggest that the glycans in GG are further elongated and larger than those in the SG.

### The glycan types are differently distributed between membrane and lumen

The glycans identified in the granule fractions may originate either from the granule membrane or the lumen (i.e., soluble protein). For SV, the lumen contains plasma proteins taken up during invagination of the membrane during neutrophil terminal differentiation ^28^. To address which of the identified glycans are displayed on membrane-associated proteins, membranes were separated from lumen for each granule population (AG, SG, GG) and the SV+PM (all obtained from three-layer gradients). Paucimannosidic glycans were enriched in the AG membranes while oligomannose glycans were enriched in the membranes of SG and SV+PM (Figure 5). In contrast, complex glycans were observed in high abundance on luminal proteins in all granule fractions.

**Figure 5.**
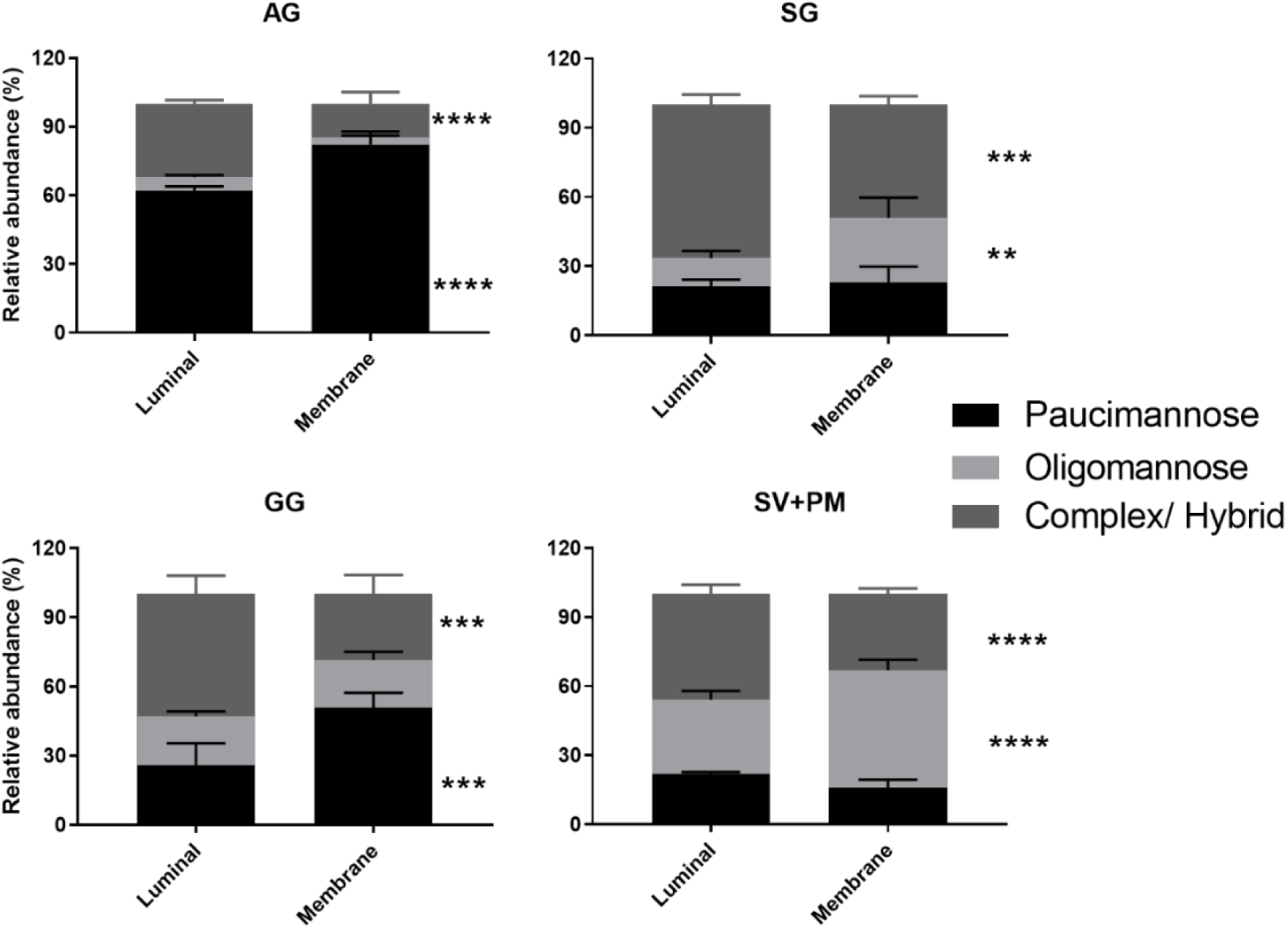
The *N*-glycan signatures differ on luminal and membrane proteins in the different neutrophil organelles. Quantitative glycomics was performed for *N*-glycans released from separate membrane and lumen (soluble) protein fractions from isolated neutrophil granules and SV+PM. The identified glycans have been grouped and quantified according to their *N*-glycan type i.e. paucimannosidic, oligomannosidic and complex/hybrid *N*-glycans. Note: Luminal glycans in SV+PM fraction are not of neutrophil origin, but comes from plasma proteins. Data represented as mean ± SEM, and unpaired two-way student’s *t*-test was used for statistical analysis. *, *p* < 0.05; **, *p* < 0.01; ***, *p* < 0.001; ****, *p* < 0.0001.

### *O*-glycans are absent in AG while extended LacNAc-rich core-2 *O*-glycans are consistently found in the other fractions

Apart from *N*-glycan analysis, proteins were also subjected to reductive β-elimination after which the released *O*-glycans were analyzed. There were no *O*-linked glycans in the AG, whereas SG+GG and SV+PM demonstrated the presence of similar *O*-glycans (Supplementary figure S5), with a total of 17 *O*-glycans identified. The most abundant *O*-glycans were mono- and di-sialylated core-2 glycans (Figure 6a, Supplementary Figure S5). Analogous to the LacNAc extension of the *N*-glycans, only one of the branches was elongated with LacNAcs on *O*-glycans. Sialic acid linked to galactose was found on the 3’ arm but not on the 6’ arm, while the fucose moiety was located on the 6’ arm together with LacNAc extensions (Figure 6b). Mono- and di-sialylated core-1 glycans were also identified at high abundance (Supplementary Figure S5). Similar to *N*-glycans, *O*-glycans with sLe^x^ epitopes were identified, with fucosylation on 6’ arm and sialylation capping on the other arm (Figure 6b). In conclusion, neutrophil granule proteins are modified with core 1 and core 2 *O*-glycans carrying LacNAc chains and Le^x^ antigens while sLe^x^ epitopes were not commonly observed.

**Figure 6.**
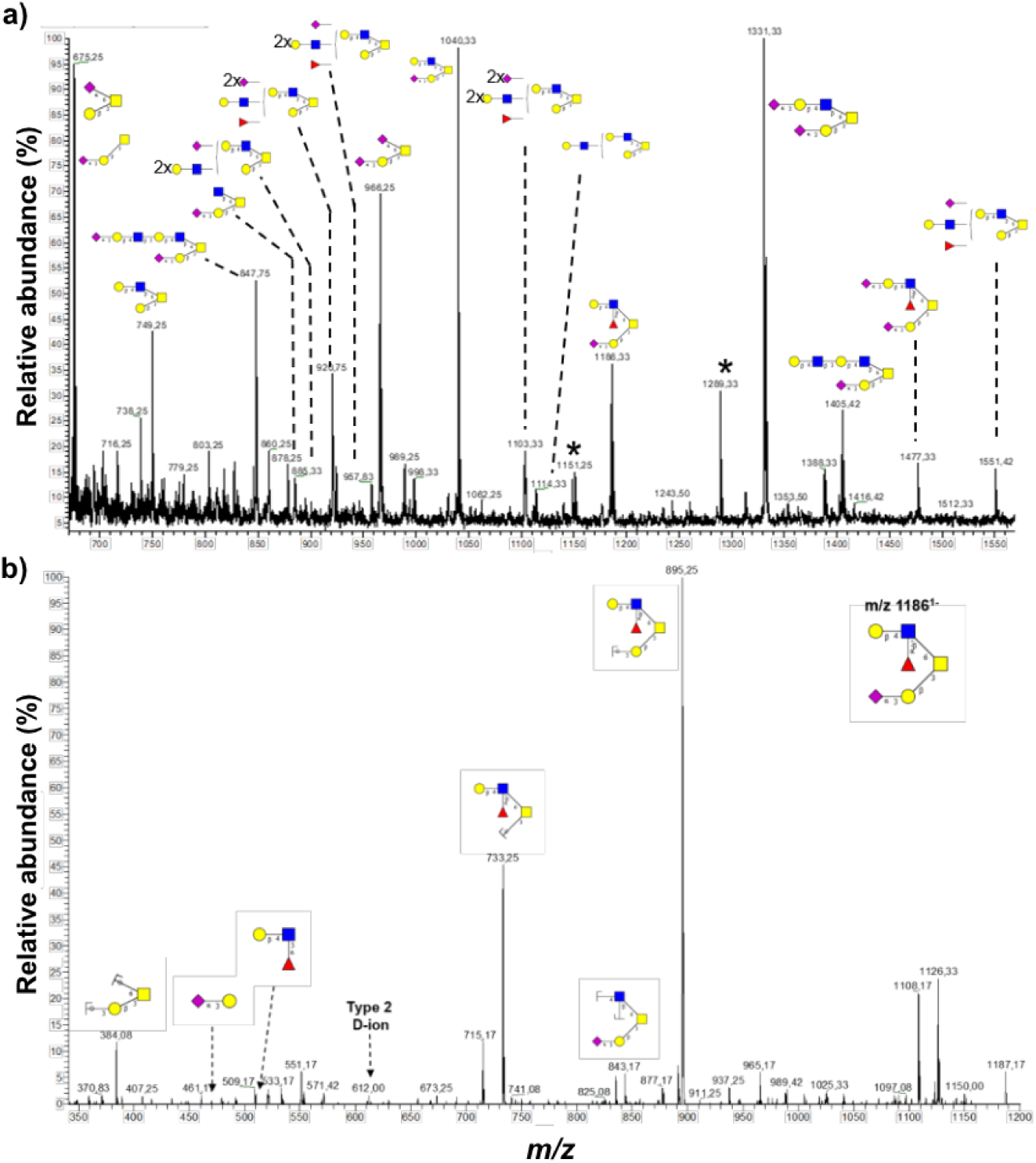
Uniform core 2 *O*-glycosylation across neutrophil granules except for the *O*-glycan-deficient Azurophil granules. a) Core 2 *O*-glycans were commonly observed with LacNAc extensions on the 6’ arm and sialic acid capping on the 3’arm galactose. b) LC-MS/MS spectra of core 2 *O*-glycan (*m/z* 1186^1-^) with sialic acid capping on 3’ arm and Lewis epitope on 6’ arm. Asterisk represents contaminating signals of unknown origin.

### Granule glycosylation during granulopoiesis

As mentioned, during granulopoiesis different granules are sequentially formed at different stages: myeloblasts or promyelocytes (AG), myelocytes/metamyelocytes (SG) and band cell stage (GG). The quantitative comparison of the glycan composition revealed a shift from paucimannosidic *N*-glycans and lack of *O*-glycans in the AG to LacNAc-containing complex glycans in the SG and elongated ditto in the GG (Figure 7). This suggests a change in glycosylation machinery over time during granulopoiesis. We therefore compared the glycomic data with the corresponding mRNA expression profiles (from the publicly available Bloodspot database ^29^) of enzymes responsible for glycan synthesis during granulopoiesis. We noticed a higher expression of mannosidase (MAN1A1) and hexosaminidase (HEXA), responsible for the synthesis of paucimannosidic glycans, in promyelocytes as compared to cells from later stages of granulopoiesis ^29^ (Supplementary Figure S6). In contrast, expression of B3GNT3, B4GalT4 and FUT3, responsible for the synthesis of LacNAc repeats and Le^x^, was higher in the later stages as compared to early maturation stage ^29^. Finally ST3Gal4, synthesizing sLe^x^, was found to be expressed predominantly during the final stages of maturation ^29^. In conclusion, the glycomic data together with database-mined mRNA expression data supports our hypothesis that a change in the glycosylation machinery occurs over time during granulopoiesis.

**Figure 7.**
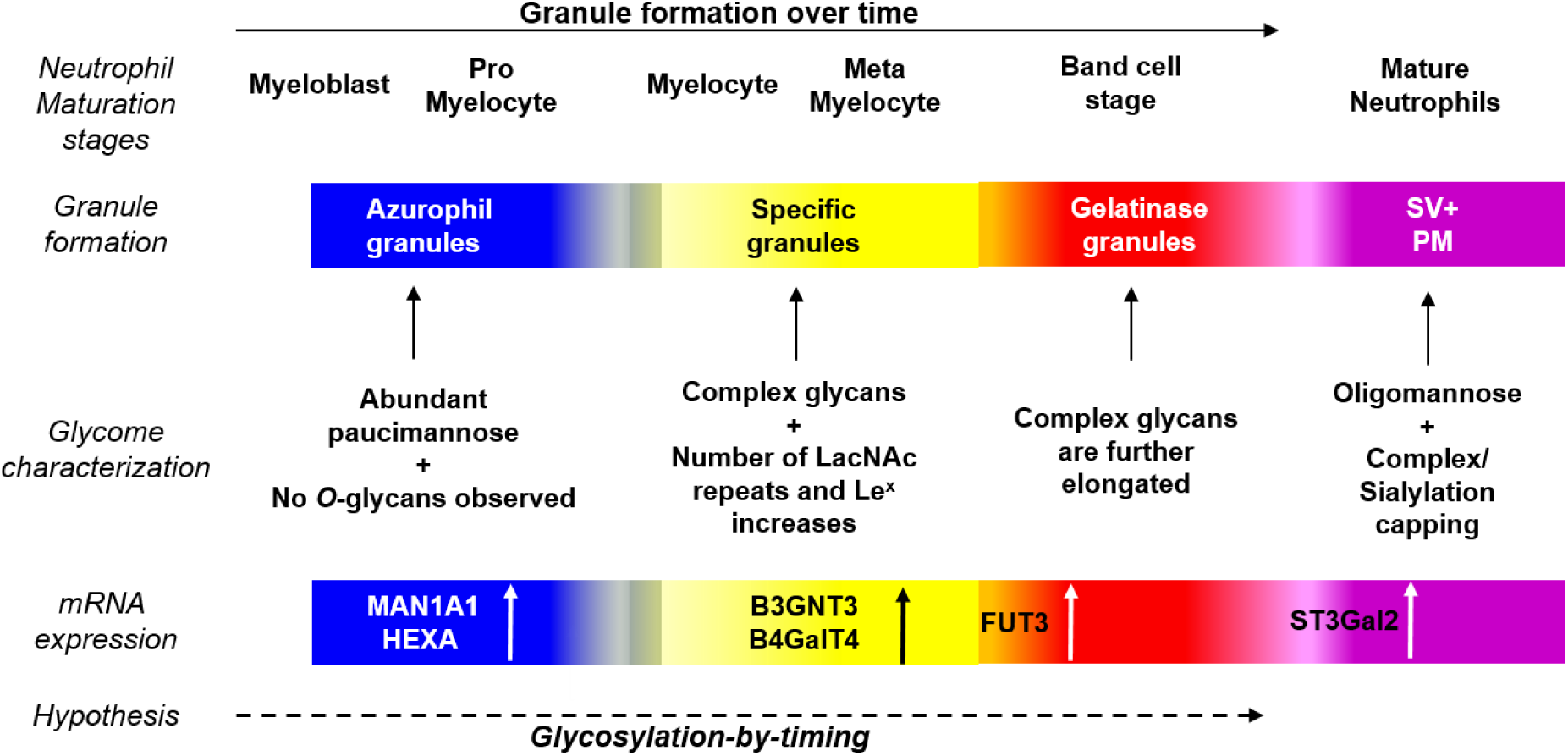
Overview of ‘glycosylation-by-timing’ hypothesis. Granule formation during neutrophil maturation is a continuum and granules carry a vast array of glycans. From the presence shorter paucimannosidic glycans in AG, to ever elongating complex glycans in SG and subsequently in GG and the corresponding mRNA expression of different glycosidases and glycosyltransferases, suggests an evolution of granule glycosylation over time. Similar to “targeting-by-timing” hypothesis describing the temporal protein synthesis of maturing neutrophils, we hypothesize a parallel ‘glycosylation-by-timing’ granule glycan synthesis that takes place during neutrophil maturation.

## Discussion

Defining molecular differences between neutrophil organelles, proteins and finally their glycosylation, is a prerequisite for better understanding neutrophil biology and function. Mapping of the protein composition of neutrophil subcellular organelles ^3,27^, as well as their gene and protein expression during granulopoiesis ^4,30,31^, has provided a comprehensive qualitative and quantitative basis for the targeting-by-timing hypothesis, which appears to accurately describe the evolution of neutrophil protein composition over time during granulopoiesis. This work provides a platform on which maturation-associated post-translational modifications, not least glycosylation, can be studied. We have, as a first, characterized the *N*- and *O*-glycans in neutrophil granules as well as in the SV/PM using a PGC-LC-MS/MS-based glycomics approach recognized for providing information of glycan structures while also giving quantitative information ^32^.

The characteristic feature of the AG glycoproteins is the presence of paucimannosidic type *N*-glycans and the absence of *O*-glycans. Our developing knowledge of protein paucimannosylation in human biology including aspects covering its structure, function, and tissue expression has recently been extensively summarized ^33^. Sequential trimming of the glycoprotein intermediate (Man3GlcNAc ± core fucose) by HexA and presumably a linkage-specific α-mannosidase gives rise to paucimannosidic glycans of which one species (M2F) represents > 60% of the total amount of paucimannose. This was supported by a strong co-localization of HexA and MPO in differentiated HL-60 cells using immunohistochemistry ^20^.

Upon inflammation *in vivo*, the AG proteins with linked paucimannosidic glycans are released or leaks from activated neutrophils as a result of substantial cell activation or cell death, carried by an ensemble of bioactive AG proteins, including MPO and serine proteases ^20,21,34,35^. The structure-function relationship of the paucimannosidic glycans, responsible for the strong proinflammatory effects of these proteins, can at this point only be speculated upon, but both paucimannosidic and oligomannosidic-type glycans have been shown to bind to mannose recognizing C-type lectin receptors expressed by macrophages and dendritic cells ^21,36^. Also, the macrophage mannose-receptor has been shown to internalize neutrophil-derived MPO^37^, playing a potential physiological role in regulating extracellular MPO levels. Although the preference of the mannose-receptor for specific glycan substrates has yet to be documented in neutrophils, the structures and localization in the various granule compartments identified here will be useful for future investigation.

Abundance of complex *N*- and *O*-glycans with extended LacNAc chains was a characteristic feature observed in the SG+GG, formed during the middle stages of granulopoiesis. The level of LacNAc chains identified are unique to neutrophils and have to our knowledge never been found in other immune cells. In fact, the number of LacNAc repeats reported here is probably underestimated, as we incidentally identified structures with up to 16 LacNacs in few of the spectra. Given that the complex glycans in the SG, GG and SV+PM fractions are mostly bi-antennary, these long LacNAc extensions placed close together on a cell surface could build a dense net for interactions with lectins, e-g., galactose-binding lectins such as galectins. In the context of neutrophil function and regulation, carbohydrate-dependent binding of galectin-3 can mediate cell adhesion, phagocytosis and activation of the respiratory burst ^38,39^. Interestingly, a prerequisite for the neutrophil response to galectin-3 is that the cells first are primed by an inflammatory stimulus and have mobilized the GG and SG, thereby having upregulated granule-stored, LacNAc-bearing protein receptors to the cell surface for interactions with galectin-3 ^40^.

The LacNac chain length may be biologically important in this context, as shown by a galectin-3 mutant carrying increased affinity for extended LacNAc structures resulting in an enhanced respiratory burst response in primed neutrophils as compared to wild-type galectin-3 ^41^. Furthermore, the LacNAc chains in the SG, GG and SV+PM were decorated with fucose and sialic acid residues, forming Lewis epitopes at the non-reducing end of *N*- and *O*-glycans. Beside the well-characterized role of Le^x^ and sLe^x^ in cell adhesion, cell priming and phagocytosis ^12,42-44^, the expression of Le^a^ epitopes on neutrophils has recently been shown to enhance neutrophil transmigration and to play a role in regulating inflammation ^45^.

Unlike AG and SG+GG, which are enriched in paucimannosidic and complex N-glycans, respectively, the SV+PM fraction contains a mix of oligomannose and processed complex glycans. While the lumen fraction of SV+PM carries proteins of non-neutrophil origin (i.e., plasma proteins invaginated into the SV ^28^), the granule membrane protein glycosylation is of neutrophil origin. The SV+PM membrane fraction was predominantly composed of oligomannose glycans that are readily available recognition tags on the cell surface of resting or slightly primed cells (having mobilized the secretory vesicles). Indeed, a historically well-known mannose-dependent neutrophil interaction is that of the bacterial mannose-binding adhesin FimH which recognizes several mannose structures on the surface of neutrophils ^46^, mediating bacterial binding and phagocytosis ^47^.

Biosynthetically, protein synthesis and granule targeting are best described by the targeting-by-timing hypothesis, where proteins synthesized at the same time during granulopoiesis end up in the same granules. How glycosylation fits into such a scheme is an interesting question with implications for cell regulation and function. Our data show distinct patterns of glycosylation for the different granules and organelles, and close analysis of these actually provide some insights; the granule-specific glycans and corresponding mRNA expression of enzymes responsible for glycan synthesis appear to co-emerge at different stages of granulopoiesis, which suggests a change in glycosylation machinery over time. The plasma membrane, which contains both unprocessed oligomannose and processed complex type glycans, is not formed within a defined time interval during neutrophil differentiation, as the granules are. This is in line with that PM proteins are successively being modified with the glycans specific to each different developmental stage during granulopoiesis, and are sorted to the cell surface. This results in the presence of oligomannose type glycans potentially produced during early stages of granulopoiesis, together with differently matured poly-LacNAc-containing glycans produced later on. The PM targeted glycans will then be endocytosed and incorporated into the membranes of the secretory vesicles during the final stages of cell maturation.

To this background, it is tempting to suggest that a protein synthesized at a certain time during granulopoiesis and targeted to a certain granule subtype, is specifically modified with a dedicated glycan, synthesized at that same time, by a glycosylation machinery that is concurrently expressed. Consequently, we have formulated a complementing hypothesis to targeting-by-timing, stating that protein glycosylation in neutrophils takes place through a “glycosylation-by-timing” process (Figure 7).

Furthermore, we suggest that the variation of glycosylation during granulopoiesis may correlate to the evolution of the Golgi complex, for which the size and number of stacks vary greatly during granulopoiesis ^48,49^, and/or the extensive glycan processing that may occur at different rates within the formed granules during neutrophil maturation, directions which we are currently pursuing. Also, the finding that the minute amounts of elastase found outside of AG and SG appear to be similarly glycosylated ^21^, may indicate that certain proteins (or glycans) do not follow glycosylation-by-timing. We are studying granular glycoproteomics to further investigate the specific protein carriers of each glycan, which will give us better understanding of how protein glycosylation is evolving.

In conclusion, we provide a map of subcellular glycans in healthy human neutrophils that displays peculiar granule-specific glycan signatures, and this first-time mapping of neutrophil granule glycans provides information on adaptations that may occur in the glycosylation machinery during granulopoiesis.

## Supporting information

Suppementary Figures

## Acknowledgements

This work was supported by the Swedish Research Council (2018-03077 and 2019-01123), the King Gustaf V 80-year Memorial Foundation, the Swedish Heart-Lung Foundation, and the Swedish government under the agreement concerning research and education of physicians (ALF). The mass spectrometer (LTQ) was supported by the Knut and Alice Wallenberg Foundation.

## Abbreviations

AG: Azurophil granules
SG: Specific granules
GG: Gelatinase granules
SV: Secretory vesicles
PM: Plasma membrane
PGC: Porous graphitized carbon
LC: Liquid chromatography
MS/MS: Tandem mass spectrometry
LacNAc: *N*-acetyl lactosamine
Le^x^: Lewis X
sLe^x^: Sialyl- Lewis X

## References

1. Borregaard N, Cowland JB. Granules of the human neutrophilic polymorphonuclear leukocyte. Blood. 1997;89(10):3503–3521.

2. Faurschou M, Borregaard N. Neutrophil granules and secretory vesicles in inflammation. Microbes Infect. 2003;5(14):1317–1327.

3. Rorvig S, Ostergaard O, Heegaard NH, Borregaard N. Proteome profiling of human neutrophil granule subsets, secretory vesicles, and cell membrane: correlation with transcriptome profiling of neutrophil precursors. J Leukoc Biol. 2013;94(4):711–721.

4. Le Cabec V, Cowland JB, Calafat J, Borregaard N. Targeting of proteins to granule subsets is determined by timing and not by sorting: The specific granule protein NGAL is localized to azurophil granules when expressed in HL-60 cells. Proc Natl Acad Sci U S A. 1996;93(13):6454–6457.

5. Phillips ML, Nudelman E, Gaeta FC, et al. ELAM-1 mediates cell adhesion by recognition of a carbohydrate ligand, sialyl-Lex. Science. 1990;250(4984):1130–1132.

6. Walz G, Aruffo A, Kolanus W, Bevilacqua M, Seed B. Recognition by ELAM-1 of the sialyl-Lex determinant on myeloid and tumor cells. Science. 1990;250(4984):1132–1135.

7. Yamaoka A, Kuwabara I, Frigeri LG, Liu FT. A human lectin, galectin-3 (epsilon bp/Mac-2), stimulates superoxide production by neutrophils. J Immunol. 1995;154(7):3479–3487.

8. Carlin AF, Uchiyama S, Chang YC, Lewis AL, Nizet V, Varki A. Molecular mimicry of host sialylated glycans allows a bacterial pathogen to engage neutrophil Siglec-9 and dampen the innate immune response. Blood. 2009;113(14):3333–3336.

9. Graham SA, Antonopoulos A, Hitchen PG, et al. Identification of neutrophil granule glycoproteins as Lewis(x)-containing ligands cleared by the scavenger receptor C-type lectin. J Biol Chem. 2011;286(27):24336–24349.

10. Babu P, North SJ, Jang-Lee J, et al. Structural characterisation of neutrophil glycans by ultra sensitive mass spectrometric glycomics methodology. Glycoconj J. 2009;26(8):975–986.

11. Karlsson A, Miller-Podraza H, Johansson P, Karlsson KA, Dahlgren C, Teneberg S. Different glycosphingolipid composition in human neutrophil subcellular compartments. Glycoconj J. 2001;18(3):231–243.

12. Foxall C, Watson SR, Dowbenko D, et al. The three members of the selectin receptor family recognize a common carbohydrate epitope, the sialyl Lewis(x) oligosaccharide. J Cell Biol. 1992;117(4):895–902.

13. Nimrichter L, Burdick MM, Aoki K, et al. E-selectin receptors on human leukocytes. Blood. 2008;112(9):3744–3752.

14. Etzioni A, Frydman M, Pollack S, et al. Brief report: recurrent severe infections caused by a novel leukocyte adhesion deficiency. N Engl J Med. 1992;327(25):1789–1792.

15. Etzioni A, Harlan JM, Pollack S, Phillips LM, Gershoni-Baruch R, Paulson JC. Leukocyte adhesion deficiency (LAD) II: a new adhesion defect due to absence of sialyl Lewis X, the ligand for selectins. Immunodeficiency. 1993;4(1-4):307–308.

16. Lucka L, Fernando M, Grunow D, et al. Identification of Lewis x structures of the cell adhesion molecule CEACAM1 from human granulocytes. Glycobiology. 2005;15(1):87–100.

17. Poland DC, Garcia Vallejo JJ, Niessen HW, et al. Activated human PMN synthesize and release a strongly fucosylated glycoform of alpha1-acid glycoprotein, which is transiently deposited in human myocardial infarction. J Leukoc Biol. 2005;78(2):453–461.

18. Theilgaard-Monch K, Jacobsen LC, Rasmussen T, et al. Highly glycosylated alpha1-acid glycoprotein is synthesized in myelocytes, stored in secondary granules, and released by activated neutrophils. J Leukoc Biol. 2005;78(2):462–470.

19. Venkatakrishnan V, Thaysen-Andersen M, Chen SC, Nevalainen H, Packer NH. Cystic fibrosis and bacterial colonization define the sputum N-glycosylation phenotype. Glycobiology. 2015;25(1):88–100.

20. Thaysen-Andersen M, Venkatakrishnan V, Loke I, et al. Human neutrophils secrete bioactive paucimannosidic proteins from azurophilic granules into pathogen-infected sputum. J Biol Chem. 2015;290(14):8789–8802.

21. Loke I, Ostergaard O, Heegaard NHH, Packer NH, Thaysen-Andersen M. Paucimannose-Rich N-glycosylation of Spatiotemporally Regulated Human Neutrophil Elastase Modulates Its Immune Functions. Mol Cell Proteomics. 2017;16(8):1507–1527.

22. Boyum A, Lovhaug D, Tresland L, Nordlie EM. Separation of leucocytes: improved cell purity by fine adjustments of gradient medium density and osmolality. Scand J Immunol. 1991;34(6):697–712.

23. Borregaard N, Heiple JM, Simons ER, Clark RA. Subcellular localization of the b-cytochrome component of the human neutrophil microbicidal oxidase: translocation during activation. J Cell Biol. 1983;97(1):52–61.

24. Feuk-Lagerstedt E, Movitz C, Pellme S, Dahlgren C, Karlsson A. Lipid raft proteome of the human neutrophil azurophil granule. Proteomics. 2007;7(2):194–205.

25. Jensen PH, Karlsson NG, Kolarich D, Packer NH. Structural analysis of N- and O-glycans released from glycoproteins. Nat Protoc. 2012;7(7):1299–1310.

26. Ceroni A, Maass K, Geyer H, Geyer R, Dell A, Haslam SM. GlycoWorkbench: a tool for the computer-assisted annotation of mass spectra of glycans. J Proteome Res. 2008;7(4):1650–1659.

27. Lominadze G, Ward RA, Klein JB, McLeish KR. Proteomic analysis of human neutrophils. Methods Mol Biol. 2006;332:343–356.

28. Borregaard N, Kjeldsen L, Rygaard K, et al. Stimulus-dependent secretion of plasma proteins from human neutrophils. J Clin Invest. 1992;90(1):86–96.

29. Bagger FO, Kinalis S, Rapin N. BloodSpot: a database of healthy and malignant haematopoiesis updated with purified and single cell mRNA sequencing profiles. Nucleic Acids Res. 2019;47(D1):D881–D885.

30. Mora-Jensen H, Jendholm J, Fossum A, Porse B, Borregaard N, Theilgaard-Monch K. Technical advance: immunophenotypical characterization of human neutrophil differentiation. J Leukoc Biol. 2011;90(3):629–634.

31. Theilgaard-Monch K, Jacobsen LC, Borup R, et al. The transcriptional program of terminal granulocytic differentiation. Blood. 2005;105(4):1785–1796.

32. Ashwood C, Lin CH, Thaysen-Andersen M, Packer NH. Discrimination of Isomers of Released N- and O-Glycans Using Diagnostic Product Ions in Negative Ion PGC-LC-ESI-MS/MS. J Am Soc Mass Spectrom. 2018;29(6):1194–1209.

33. Tjondro HC, Loke I, Chatterjee S, Thaysen-Andersen M. Human protein paucimannosylation: cues from the eukaryotic kingdoms. Biol Rev Camb Philos Soc. 2019;94(6):2068–2100.

34. Loke I, Packer NH, Thaysen-Andersen M. Complementary LC-MS/MS-Based N-Glycan, N-Glycopeptide, and Intact N-Glycoprotein Profiling Reveals Unconventional Asn71-Glycosylation of Human Neutrophil Cathepsin G. Biomolecules. 2015;5(3):1832–1854.

35. Reiding KR, Franc V, Huitema MG, Brouwer E, Heeringa P, Heck AJR. Neutrophil myeloperoxidase harbors distinct site-specific peculiarities in its glycosylation. J Biol Chem. 2019;294(52):20233–20245.

36. Feinberg H, Mitchell DA, Drickamer K, Weis WI. Structural basis for selective recognition of oligosaccharides by DC-SIGN and DC-SIGNR. Science. 2001;294(5549):2163–2166.

37. Shepherd VL, Hoidal JR. Clearance of neutrophil-derived myeloperoxidase by the macrophage mannose receptor. Am J Respir Cell Mol Biol. 1990;2(4):335–340.

38. Karlsson A, Christenson K, Matlak M, et al. Galectin-3 functions as an opsonin and enhances the macrophage clearance of apoptotic neutrophils. Glycobiology. 2009;19(1):16–20.

39. Kuwabara I, Liu FT. Galectin-3 promotes adhesion of human neutrophils to laminin. J Immunol. 1996;156(10):3939–3944.

40. Feuk-Lagerstedt E, Jordan ET, Leffler H, Dahlgren C, Karlsson A. Identification of CD66a and CD66b as the major galectin-3 receptor candidates in human neutrophils. J Immunol. 1999;163(10):5592–5598.

41. Salomonsson E, Carlsson MC, Osla V, et al. Mutational tuning of galectin-3 specificity and biological function. J Biol Chem. 2010;285(45):35079–35091.

42. Polley MJ, Phillips ML, Wayner E, et al. CD62 and endothelial cell-leukocyte adhesion molecule 1 (ELAM-1) recognize the same carbohydrate ligand, sialyl-Lewis x. Proc Natl Acad Sci U S A. 1991;88(14):6224–6228.

43. Brazil JC, Sumagin R, Cummings RD, Louis NA, Parkos CA. Targeting of Neutrophil Lewis X Blocks Transepithelial Migration and Increases Phagocytosis and Degranulation. Am J Pathol. 2016;186(2):297–311.

44. Beauharnois ME, Lindquist KC, Marathe D, et al. Affinity and kinetics of sialyl Lewis-X and core-2 based oligosaccharides binding to L- and P-selectin. Biochemistry. 2005;44(27):9507–9519.

45. Brazil JC, Sumagin R, Stowell SR, et al. Expression of Lewis-a glycans on polymorphonuclear leukocytes augments function by increasing transmigration. J Leukoc Biol. 2017;102(3):753–762.

46. Sauer MM, Jakob RP, Luber T, et al. Binding of the bacterial adhesin FimH to its natural, multivalent high-mannose type glycan targets. J Am Chem Soc. 2018.

47. Tewari R, MacGregor JI, Ikeda T, Little JR, Hultgren SJ, Abraham SN. Neutrophil activation by nascent FimH subunits of type 1 fimbriae purified from the periplasm of Escherichia coli. J Biol Chem. 1993;268(4):3009–3015.

48. Bainton DF, Farquhar MG. Origin of granules in polymorphonuclear leukocytes. Two types derived from opposite faces of the Golgi complex in developing granulocytes. J Cell Biol. 1966;28(2):277–301.

49. Bainton DF, Ullyot JL, Farquhar MG. The development of neutrophilic polymorphonuclear leukocytes in human bone marrow. J Exp Med. 1971;134(4):907–934.

