## Supplementary material for "Glycan analysis of human neutrophil granules implicates a maturation-dependent glycosylation machinery": Suppementary Figures

### These authors contributed equally

***Corresponding author**

**Short title**: Glycomic characterization of neutrophil granules

#### Supplementary Figure S1

Representative mass spectral profile of AG N-glycans illustrating the presence of the six paucimannosidic glycans (*m/z* 587.2$^{1-}$, 733.2$^{1-}$, 749.2$^{1-}$, 895.3$^{1-}$, 911.3$^{1-}$ and 1057.3$^{1-}$) and the most abundant complex *N*-glycan (*m/z* 856.3$^{2-}$).

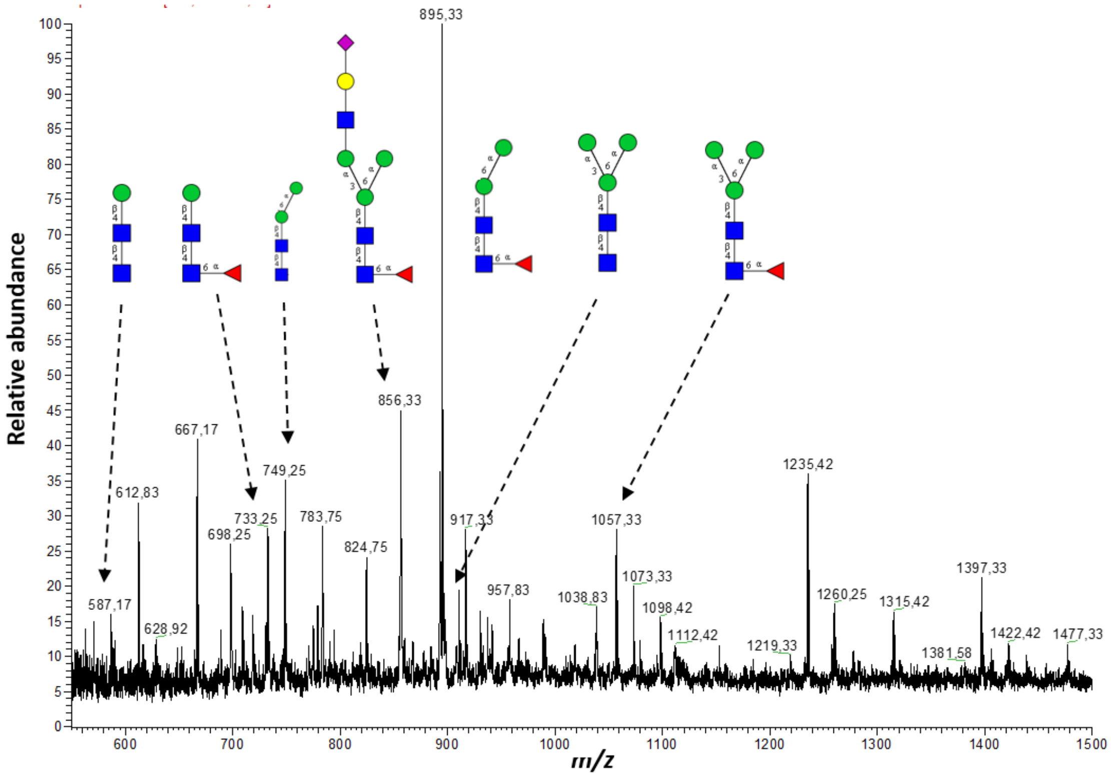

**Supplementary Figure S2**

Representative mass spectrum of SV+PM fraction showing the presence of the abundant high mannose (marked in boxes) and the high *m/z* region illustrating the elongated complex glycans as observed in SG+GG fraction.

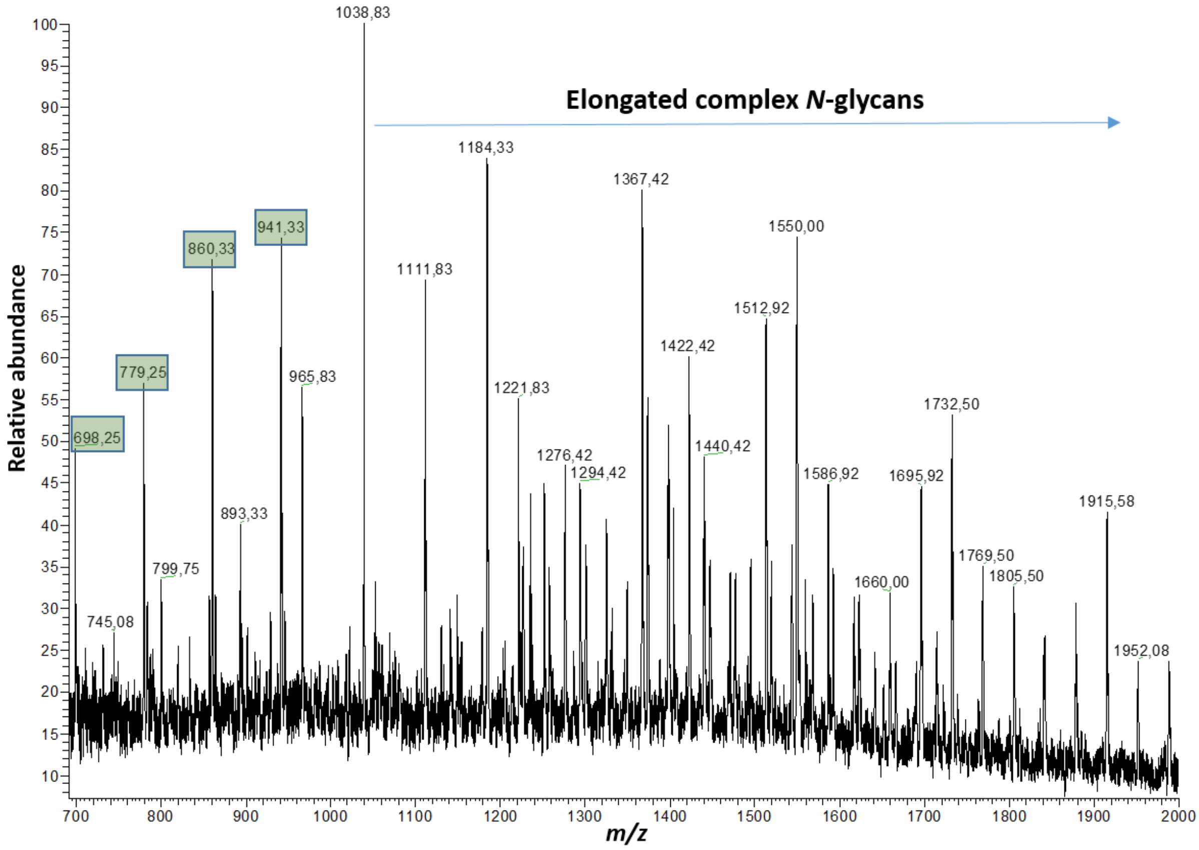

#### Supplementary Figure S3

MS/MS fragmentation spectrum of *m/z* 1367 corresponding to an *N*-glycan containing three LacNAc units with fucose and sialic acid, showing the presence of LacNAc repeats on the 6' arm.

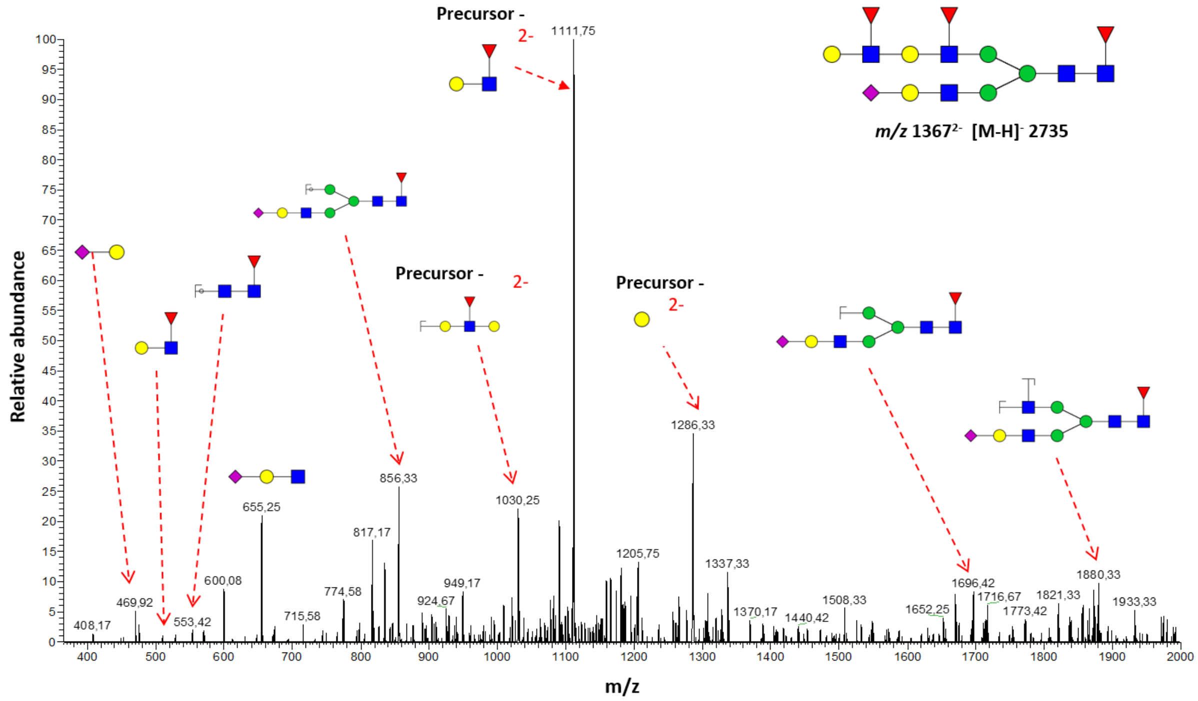

**Supplementary Figure S4**

Figure shows the relative MS intensities (in %) of structures containing LacNAcs (3 to 10) and d) Lewis epitopes (0 to 7), which are identified in the SG+GG and SV+PM fraction, respectively.

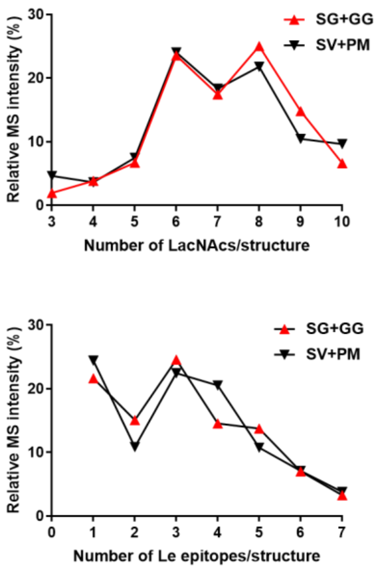

**Supplementary Figure S5**

Mass spectral profile of O-glycans analyzed from SV+PM fraction, illustrating i) abundant O-glycans identified and ii) the similarity between SG+GG and SV+PM O-glycosylation. Some of the abundant O-glycans identified are represented.

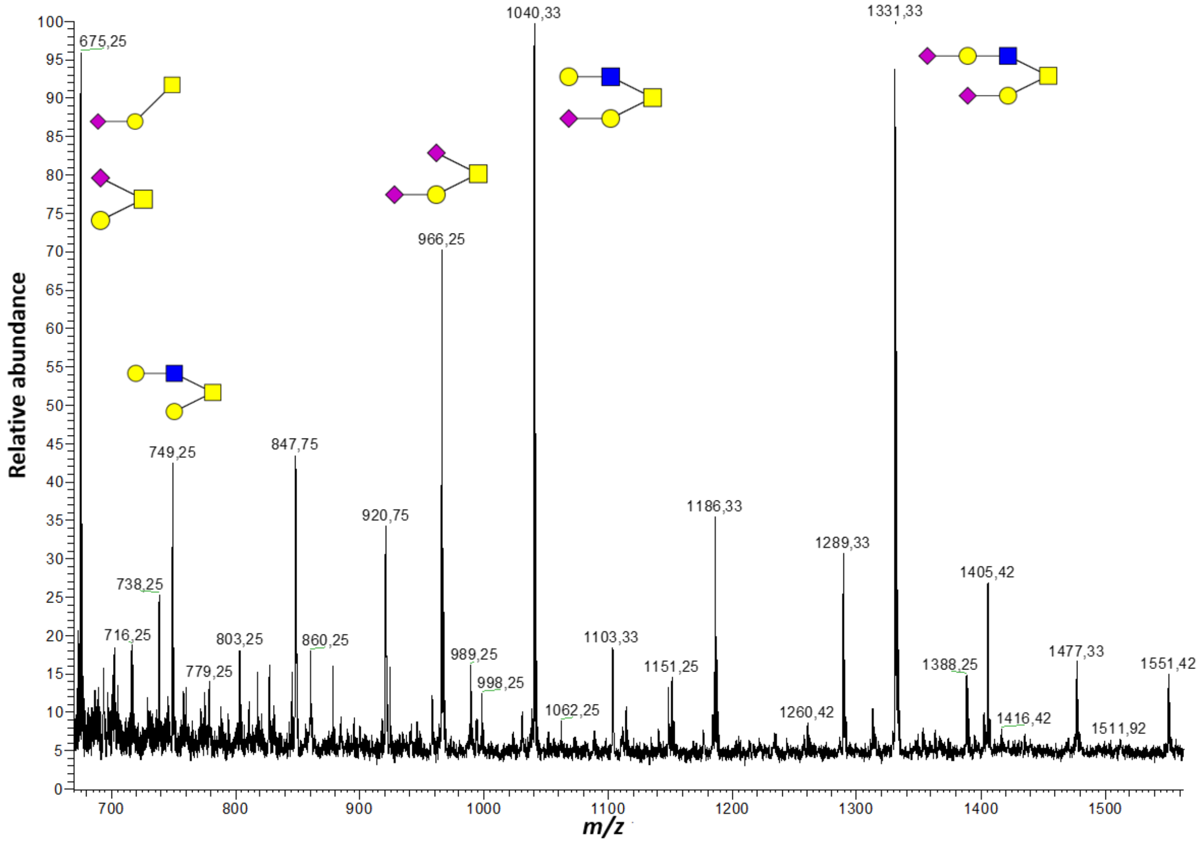

**Supplementary Figure S6**

The mRNA expression profiles of various glycosidases and glycosyl transferases at different stages during neutrophil maturation in bone marrow. The different stages are referred as MB/PM (myeloblast/promyleocyte), MY/MM (myelocyte/metamyelocyte) and BC/PMN (band cell stae/mature PMN). The data represented here were obtained from publicly available database “Bloodspot”.

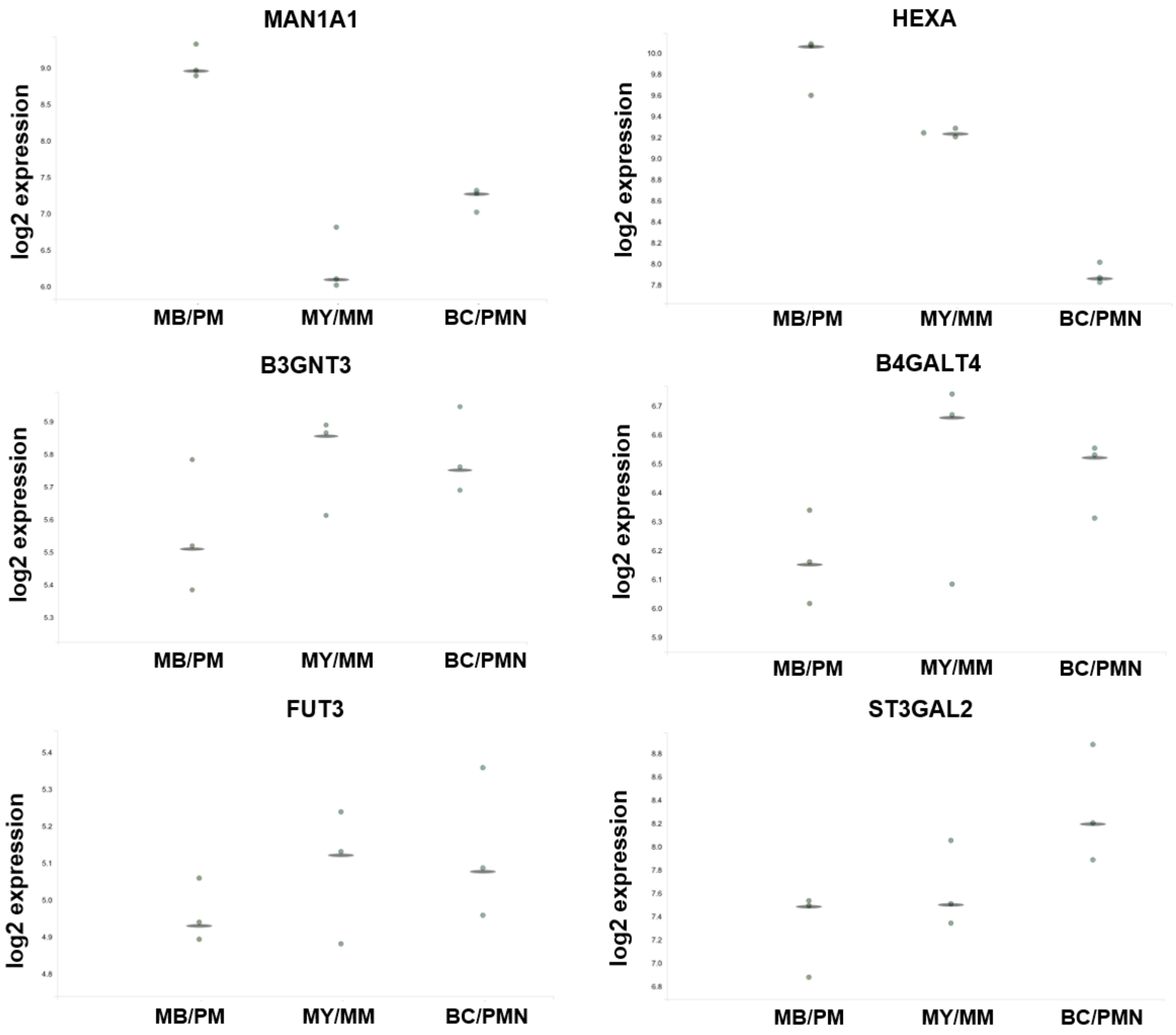
